# A conserved extracellular ribonuclease with broad-spectrum cytotoxic activity enables smut fungi to compete with host-associated bacteria

**DOI:** 10.1101/2023.04.19.537453

**Authors:** Bilal Ökmen, Philipp Katzy, Raphael Wemhöner, Gunther Doehlemann

## Abstract

Ribotoxins are secreted ribonucleases that specifically target and cleave the universally conserved sarcin-ricin loop sequence of rRNA, which leads to inhibition of protein biosynthesis and subsequently to cell death. We have identified and characterized a secreted Ribo1 protein of plant pathogenic smut fungi. Heterologous expression in different model systems showed that smut Ribo1 has cytotoxic activity against bacteria, yeast, host and non-host plants. Recombinant expression of Ribo1 in *Nicotiana benthamiana* induced plant cell death; however, an active site mutant induced cell death only when expressed as a secreted protein. In the maize smut *Ustilago maydis*, transcription of *Ribo1* is specifically induced in early infection stages. While a knock-out mutant revealed that Ribo1 is dispensable for *U. maydis* virulence, the overexpression of Ribo1 *in-planta* had a strong dominant negative effect on virulence and induced host defense responses including cell death. This suggests a function of Ribo1 during the epiphytic development rather than for invasive colonization of the host. Accordingly, in presence of the biocontrol bacteria *Pantoea* sp., which were isolated from maize leaves, the *ribo1* knock-out mutant was significantly impaired in virulence. Together, we conclude that Ribo1 enables smut fungi to compete with host-associated bacteria during epiphytic development.

## Introduction

Plants have a lifelong interaction with a complex and diverse microbial community that inhabits both their underground and aboveground organs, which is collectively called the plant microbiota. Plant associated microbial communities can support growth of their host by supplying nutrients, increasing tolerance to both biotic and abiotic stresses (Hiruma *et al*., 2016; Almario *et al*., 2017; Vorholt *et al*., 2017; Harbort *et al*., 2020). In plant-pathogen interactions, however, microbial pathogens reduce the fitness of their host plants, which is a major threat to global food production. During host colonization, plant pathogens deploy adapted repertoires of effector proteins, which interfere with host defense and physiology to promote disease. Effectors have wide range of functions, such as the inhibition of host defense related enzymes, hydrolyzation and/or detoxification of antimicrobial compounds, modulation of host metabolic pathways, or the protection of fungal cell walls against lytic enzymes of the host plant (Ökmen & Doehlemann, 2014; Toruño *et al*., 2016). Recent studies revealed the importance of host associated bacteria for plant survival and protection against phytopathogenic fungi and oomycetes (Cha *et al*., 2016; Durán *et al*., 2018). For example, Carrión *et al*., (2019) reported that upon pathogen invasion, members of *Chitinophaga* and *Flavobacterium* bacterial species are enriched in the endosphere of the host and enhance enzymatic activity of fungal cell wall degrading enzymes, which results in protection against fungal infection (Carrión *et al*., 2019). For this reason, the plant microbiota can be seen as an external shell providing an additional line of defense against pathogens. Therefore, besides modulating host physiological processes, phytopathogens also need to modulate host microbiota for successful colonization. Recently, several studies have reported the role of effectors in the modulation of host microbial communities (Kettles, Graeme J. *et al*., 2018; Snelders *et al*., 2018; Snelders, N *et al*., 2021).

Smut fungi are non-obligate biotrophic fungal pathogens that can infect several economically important crops, such as maize, wheat, barley, oat and sugar cane (Zuo *et al*., 2019). Most of the known smut fungi penetrate the coleoptile of germinated seeds to start their pathogenic life cycle. Penetration is followed by a long phase of symptomless infection which is characterized by both extra-and intracellular proliferation of the fungus in the inflorescence of the respective host. However, only at the late stage of disease symptoms manifest as masses of dark brown smut teliospores (Zuo *et al*., 2019). Bioinformatic and transcriptomic analysis revealed the effector catalogue of the maize smuts *Ustilago maydis* (Lanver *et al*., 2018) and *Sporisorium reilianum (Zuo et al*., *2021)*, as well as of the barley smut *Ustilago hordei* (Ökmen *et al*., 2018b). In line with their biotrophic life style, smut fungi have a reduced number of plant cell wall degrading enzymes (Ökmen *et al*., 2018b; Schuster *et al*., 2018). An effector candidate that caught our attention in the effectome of smut fungi has similarity to fungal RNase T1 proteins, which belong to a large family of secreted ribonucleases. Extracellular ribotoxins stand out among the members of this family due to their cytotoxic activity. Once translocated into the host cell, ribotoxins specifically target and cleave conserved sarcin/ricin loop (SRL) of ribosomal RNA (rRNA), which leads to inhibition of protein biosynthesis and subsequently to cell death (Garcia-Ortega *et al*., 2002; García-Mayoral *et al*., 2005; Citores *et al*., 2018). For example, α-sarcin, a ribotoxin from *Aspergillus* sp., displays a strong cytotoxic activity against insects and fungal cells (Citores *et al*., 2018). In this study, we have functionally characterized a secreted ribotoxin (Ribo1) of smut fungi, with regard to its role in host-pathogen interactions. We showed that Ribo1 is exclusively expressed only at very early time point of infection and has a wide range of cytotoxic activity against bacteria and yeast cells. Our observations revealed that Ribo1 enables smut fungi to compete with the host-associated bacteria.

## Results

### Conserved smut Ribo1 induces cell death in plant

Pfam domain analysis of the *Ustilago hordei* proteome identified two proteins with a predicted ribonuclease T1 (RNase T1) domain, *UHOR_02675* and *UHOR_03174*. Both proteins contain a predicted N-terminal signal peptide (16 amino acids) for extracellular secretion, and their RNase T1 domain is characterized by a tyrosine–aspartic acid–glutamic acid-histidine (YEDH) catalytic site at amino acid positions 62, 80, 98 and 115, respectively **(Fig. 1a)**. Phylogenetic tree analysis performed with *U. hordei* RNase T1 homologs, including well known bacterial and fungal ribotoxins, showed that smut RNase T1 proteins are clustered with non-toxic fungal RNase T1 proteins **(Fig. 1b)**. The recently identified RALPHs avirulence proteins from *Blumeria graminis* f. sp. *hordei* have structural similarity to RNase T1, but appeared to be outliers in the constructed phylogenetic tree **(Fig. 1b)** (Praz *et al*., 2017).

**Fig. 1.**
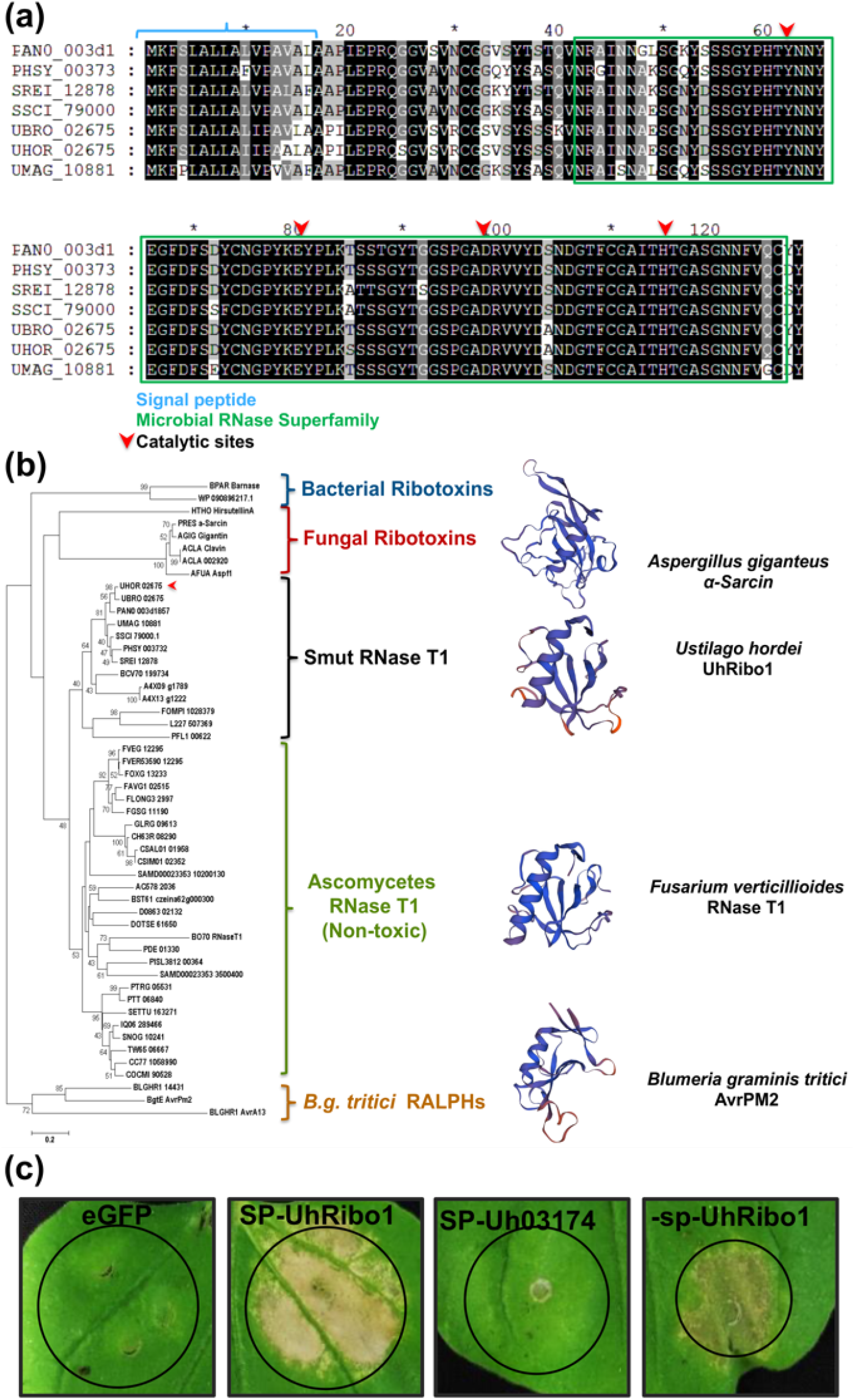
RNase T1 ribonucleases of smut fungi. **(a) Alignment of smut Ribo1**. PANO: *Moesziomyces antarcticus*; PHSY: *Pseudozyma hubeiensis* SY62; SSCI: *Sporisorium scitamineum*; SREI: *Sporisorium reilianum;* UBRO: *Ustilago bromivora*; UHOR: *Ustilago hordei*; UMAG: *Ustilago maydis*. **(b) Phylogenetic tree analysis of smut RNase T1**. A minimum evolution tree was constructed by using an alignment of the full-length amino acid sequence of RNase T1 homologs from different fungi and bacteria by using ClustalOmega. The tree was constructed by using Mega7 software by using the minimum evolution algorithm performing 1000 bootstraps. The scale bar represents the number of substitutions per site. Predicted 3D protein models were constructed by using AlphaFold protein structure database. A4X13: *Tilletia indica*; A4X09: *Tilletia walkeri*; ACLA: *Aspergillus clavatus*; AC578: *Mycosphaerella eumusae*; AFUA: *Aspergillus fumigatus*; AGIG: *Aspergillus giganteus*; BCV70DRAF: *Testicularia cyperi*; Bgt: *Blumeria graminis* f. sp. *tritici*; BLGHR: *Blumeria graminis* f. sp. *hordei*; BO70DRAFT: *Aspergillus heteromorphus*; BST61: *Cercospora zeina*; CC77DRAFT: *Alternaria alternata*; CH63R: *Colletotrichum higginsianum*: IMI: *Bipolaris oryzae*; ATCC: *Colletotrichum salicis*; CSIM: *Colletotrichum simmondsii*; D0863: *Hortaea werneckii*; DOTSE: *Dothistroma septosporum*; NZE10FAVG1: *Fusarium avenaceum*; FGSG: *Fusarium graminearum*; PH-1FLONG3: *Fusarium longipes*; FOXG: *Fusarium oxysporum* f. sp. *lycopersici*; FVER: *Fusarium verticillioides*; FOMPI: *Fomitopsis pinicola*; SS1GLRG: *Colletotrichum graminicola*; M1.001IQ06: S*tagonospora* sp.; SRC1lsM3aL227: *Lentinus tigrinus*; ALCF2SS1-6PANO: *Moesziomyces antarcticus*; PDE: *Penicillium oxalicum*; 114-2PFL1: *Anthracocystis flocculosa*; PF-1PHSY: *Pseudozyma hubeiensis*; SY62PISL3812: *Talaromyces islandicus*; PRES: *Penicillium resedanum*; PTRG: *Pyrenophora tritici-repentis*; Pt-1C-BFPPTT: *Pyrenophora teres* f. *teres*; SAMD00023353: *Rosellinia necatrix*; SC10_B2orf05621: *Bacillus paralicheniformis*; SETTU: *Exserohilum turcica*; Et28ASNOG: *Parastagonospora nodorum*; SN15SSCI: *Sporisorium scitamineum*; SREI: *Sporisorium reilianum*; SRZ2TW65: *Stemphylium lycopersici*; UBRO: *Ustilago bromivora*; UHOR: *Ustilago hordei*; UMAG: *Ustilago maydis*; WP_090896217.1: *Paenibacillus* sp. **(c) Heterologous expression of *Ribonuclease T1* (*Ribo1*) in *Nicotiana benthamiana***. By using the *Agrobacterium tumefaciens-*mediated transient expression assay, *Ustilago hordei* (±SP) *UHOR_02675, UHOR_03174* and *eGFP* were expressed in *N. benthamiana* leaves under the control of the *35SCaMV* promoter. Pictures were taken at 5 days post infiltration (dpi). White areas on the leaf surface are indicative of cell death.

To confirm that *UHOR_02675* and *UHOR_03174* are non-toxic RNases, we transiently expressed both genes in *Nicotiana benthamiana* under the control of *35S CaMV* promoter. Transient expression of the GFP-encoding gene served as negative control. The *Agrobacterium tumefaciens*-mediated expression of neither the GFP control nor the UHOR_03174 resulted in cell death induction in tobacco leaves **(Fig. 1c)**. On the contrary, the UHOR-02675 protein induced plant cell death in tobacco leaves, either when expressed with signal peptide for secretion or without **(Fig. 1c, Fig. 2a)**. Because of this cell death inducing activity, we named UHOR_02675 as UhRibo1 (*U. hordei* Ribotoxin 1). Similar to UhRibo1, heterologous expression of the *RNase T1* genes from *Ustilago maydis, Sporisorium reilianum* and *Fusarium verticillioides* also induced plant cell death in tobacco leaves after 3-5 days post infiltration (dpi), irrespective of the presence of the secretion signals **(Fig. 2a)**. To get more insight to the mode of Ribo1-induced cell death, an active site mutant of UhRibo1 (UhRibo1-M^E80Q/D98N^) was generated and expressed as C-terminally HA-tagged protein with or without signal peptide in wild type **(Fig. 2b)** and Δ*sobir1 N. benthamiana* **(Fig. S1)**.

**Fig. 2.**
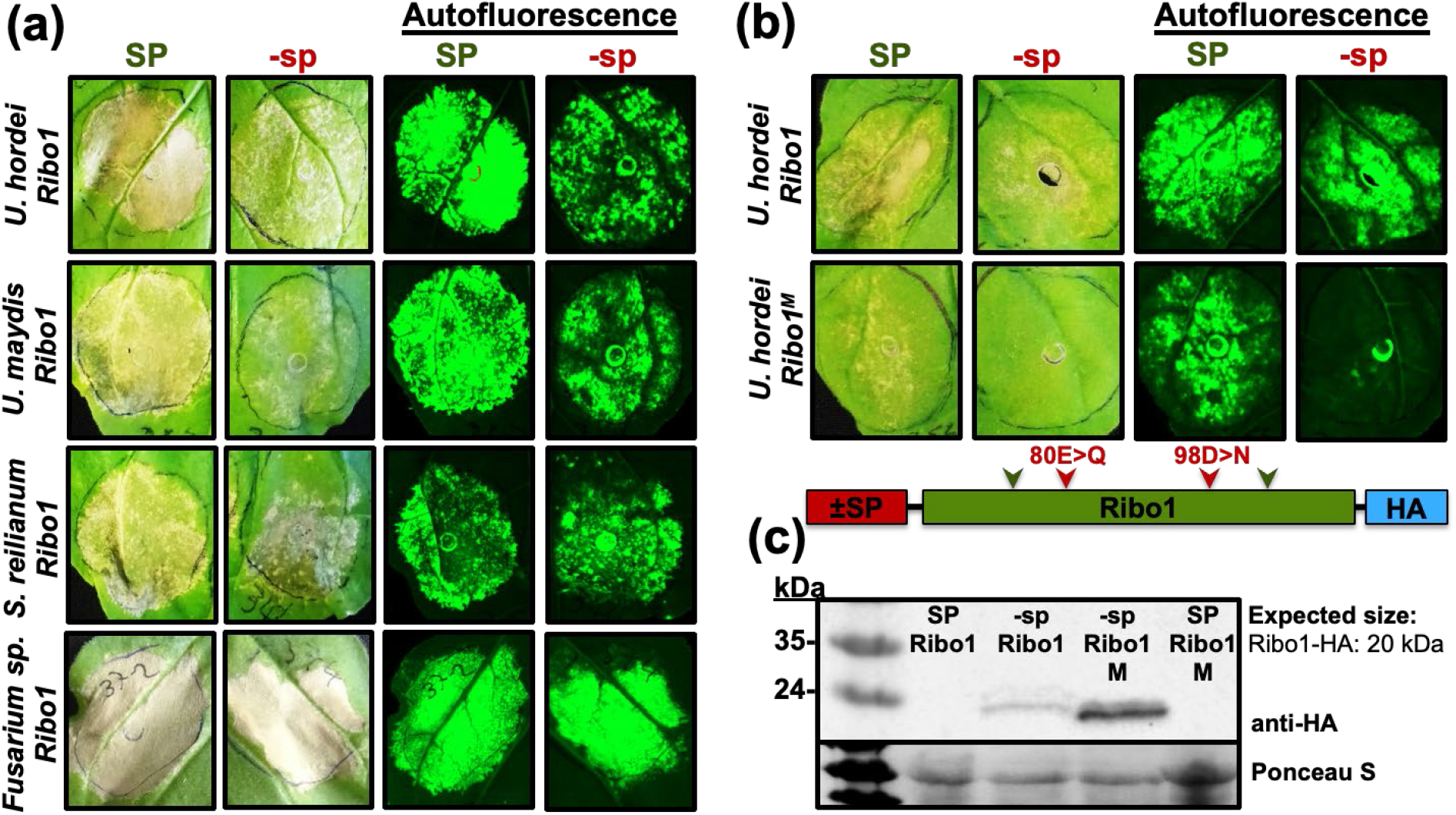
Heterologous expression of *Ribonuclease T1* (*Ribo1*) orthologs in tobacco. **(a)** By using the *A. tumefaciens-*mediated transient expression assay, *Ustilago hordei, Ustilago maydis, Sporisorium reilianum*, and *Fusarium verticillioides Ribo1* and active site mutant of *UhRibo1* were transiently expressed in *Nicotiana benthamiana* leaves under the control of *35SCaMV* promoter and with or without plant signal peptide (±SP) for extracellular secretion. Expression of *UhRibo1, UmRibo1, SrRibo1* and FvRibo1 with or without signal peptide results in plant cell death in tobacco (necrotic regions). Picture were taken after 5 days post infiltration (dpi). Autofluorescence pictures were taken with Bio-Rad Chemidoc imaging system. **(b) Heterologous expression of active site mutant version of *UhRibo1* in tobacco**. Wild type and active site mutant of *UhRibo1* were transiently expressed in *Nicotiana benthamiana* leaves under the control of *35SCaMV* promoter and with or without plant signal peptide (±SP). While secreted wild type (+SP), non-secreted *UhRibo1* (-sp) and secreted active site mutant version of UhRibo1 (Ribo1^M^) induce plant cell death, non-secreted active site mutant Ribo1^M^ did not induce cell death. Picture were taken after 5 dpi. Autofluorescence pictures were taken with Bio-Rad Chemidoc imaging system. All pictures show representative plants of at least three biological replicates. **(c)** Western blot analysis showing expression of non-secreted active site mutant version of UhRibo1. Western blot was performed with anti-HA antibody. Ponceau S staining was used to show loading control.

Suppressor of BIR1-1 (SOBIR1) is a receptor like kinase that is required for immune signaling of some plasma membrane localized receptors (Huang *et al*., 2021). C-terminally HA-tagged UhRibo1 (with and without signal peptide) was used as positive control for induction of cell death **(Fig. 2b)**. In *Agrobacterium*-mediated transient gene expression assays, extracellularly expressed UhRibo1-M^E80Q/D98N^ induced cell death in both wild type and Δ*sobir1* tobacco **(Fig. 2b** and **Fig. S1)**, while intracellular expression of UhRibo1-M^E80Q/D98N^ did not induce any cell-death **(Fig. 2b** and **Fig. S1)**. On the contrary, once again wild type UhRibo1 induced plant cell death either with or without signal peptide **(Fig. 2b** and **Fig. S1)**. Western blot analysis of total protein extracts from infiltrated leaf areas confirmed effective protein production for both wild type and active site mutant versions of UhRibo1 without secretion signal **(Fig. 2c)**. Despite the induction of plant cell death, no secreted UhRibo1 protein was detected, both for wild type and active site mutant **(Fig. 2c)**. A likely explanation for this is that the C-terminal HA-tag, which was used for detection, is cleaved after secretion to the apoplastic space. Such cleavage of affinity tags in the *N. benthamiana* apoplast has been shown in a previous study (van Esse *et al*., 2006). Together, our results suggest that the enzymatic activity of UhRibo1 is not required for the cell death induction by the secreted protein, while the enzyme activity is essential for the cell death induction by non-secreted protein.

During the cloning of *FvRibo1* from cDNA, amplification with gene specific primers revealed two splicing variants (**Fig. 3a-b**; L: long version, S: short version). Transient expression of FvRibo1_L and FvRibo1_S proteins in *N. benthamiana* showed that FvRibo1_L induced plant cell death with or without signal peptide, while FvRibo1_S did not trigger any cell death **(Fig. 3c)**. Alignment of UhRibo1, FvRibo1_L and FvRibo1_S proteins identified a stretch of 11 conserved amino acids that was absent in FvRibo1_S **(Fig. 3a)**. Deletion of this 11 aa motif from UhRibo1 completely abolished its cell death inducing activity **(Fig. 3c)**, suggesting that this sequence motif is required for the cell death inducing activity of Ribo1.

**Fig. 3.**
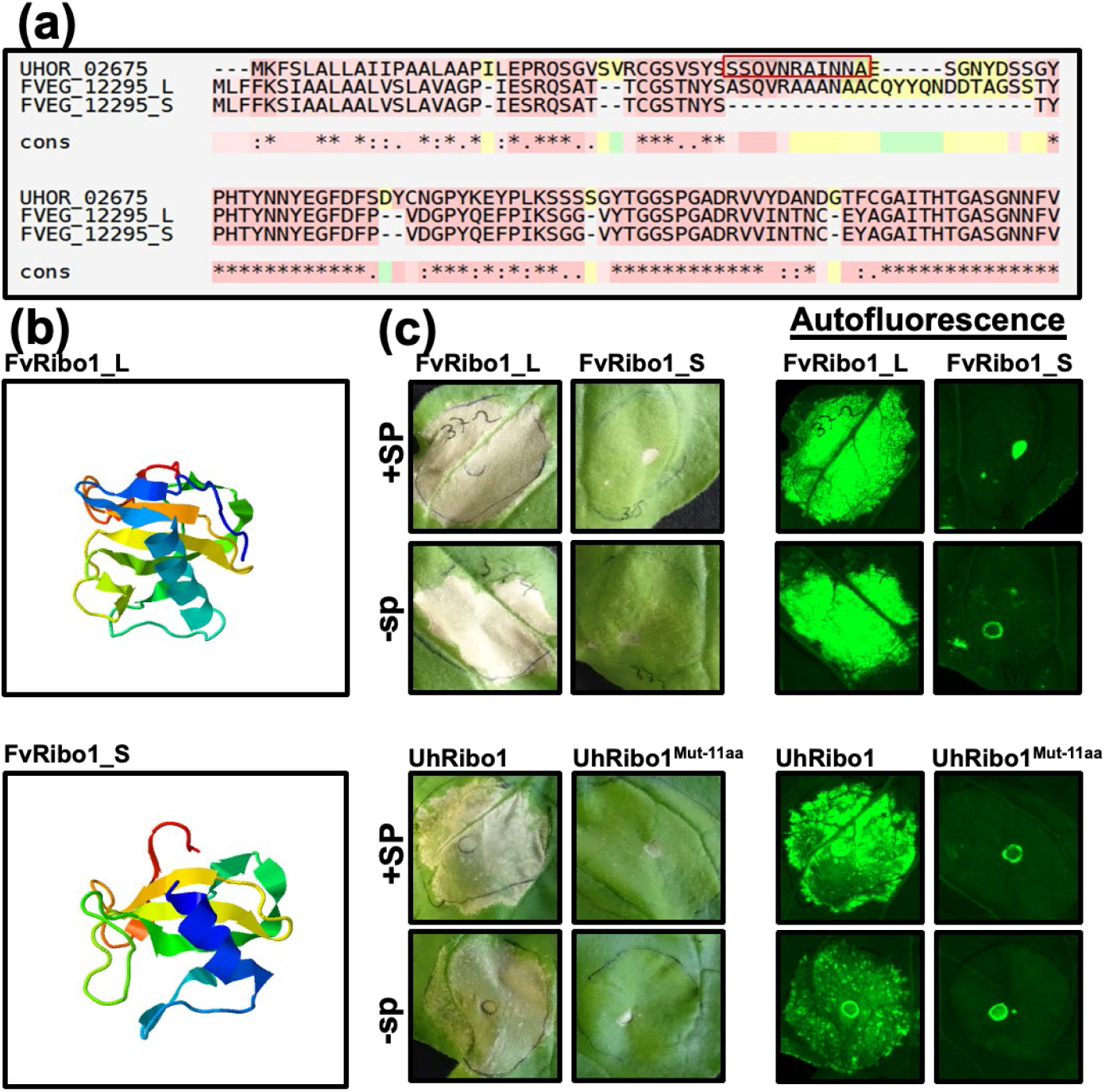
(a) Amino acid alignment of UhRibo1, long and short version of FvRibo1. Red box indicates 11 amino acids from UhRibo1 that are absent in short version of FvRibo1. **(b) 3D-modeling of long and short version of FvRibo1. (c) Deletion of 11 aa in UhRibo1 results in loss of cell-death inducing activity**. ±SP FvRibo1_L (long version), ±SP FvRibo1_S (short version), ±SP UhRibo1 and ±SP UhRibo1^Mut-11 aa^ were transiently expressed in *Nicotiana benthamiana* leaves under the control of *35SCaMV* promoter. Picture were taken after 5 days post infiltration (dpi). Autofluorescence images were taken with a Bio-Rad Chemidoc imaging system. All pictures show representative plants of at least three biological replicates.

### Smut Ribo1 is a secreted protein

To confirm secretion of Ribo1 during host colonization, Ribo1 was expressed as a secreted, C-terminally mCherry-tagged protein in the *U. maydis* strain SG200 under the control of *Umpit2* promoter, which ensures high expression levels during host colonization (Mueller *et al*., 2013). The *U. maydis-*maize pathosystem was used for characterization of Ribo1 in smut fungi, since the phenotypic observation and scoring during host colonization is much easier compared to other smuts, and the feasibility of a southern blot analysis for integration number of gene, unlike *U. hordei*. Secreted Pit2-mCherry was used as a positive control for secretion (Mueller *et al*., 2013) and internally expressed mCherry (Int.mCherry) under the control of *Umpit2* promoter was used as a negative control. Confocal microscopy of maize leaves inoculated with SG200-Ribo1-mCherry and Pit2-mCherry revealed the presence of fluorescence signals on the outer surface of fungal hyphal tips, which indicates secretion of both proteins **(Fig. 4)**. On the contrary, intracellular mCherry was detected only inside the fungal cell **(Fig. 4**). Moreover, plasmolysis with 1 M sodium chloride solution resulted in an accumulation of Ribo1-mCherry and Pit2-mCherry signals in the apoplastic space (depicted with white asterisk) **(Fig. 4)**. From this result, we conclude that Ribo1 protein is secreted from *U. maydis* hyphae.

**Fig. 4.**
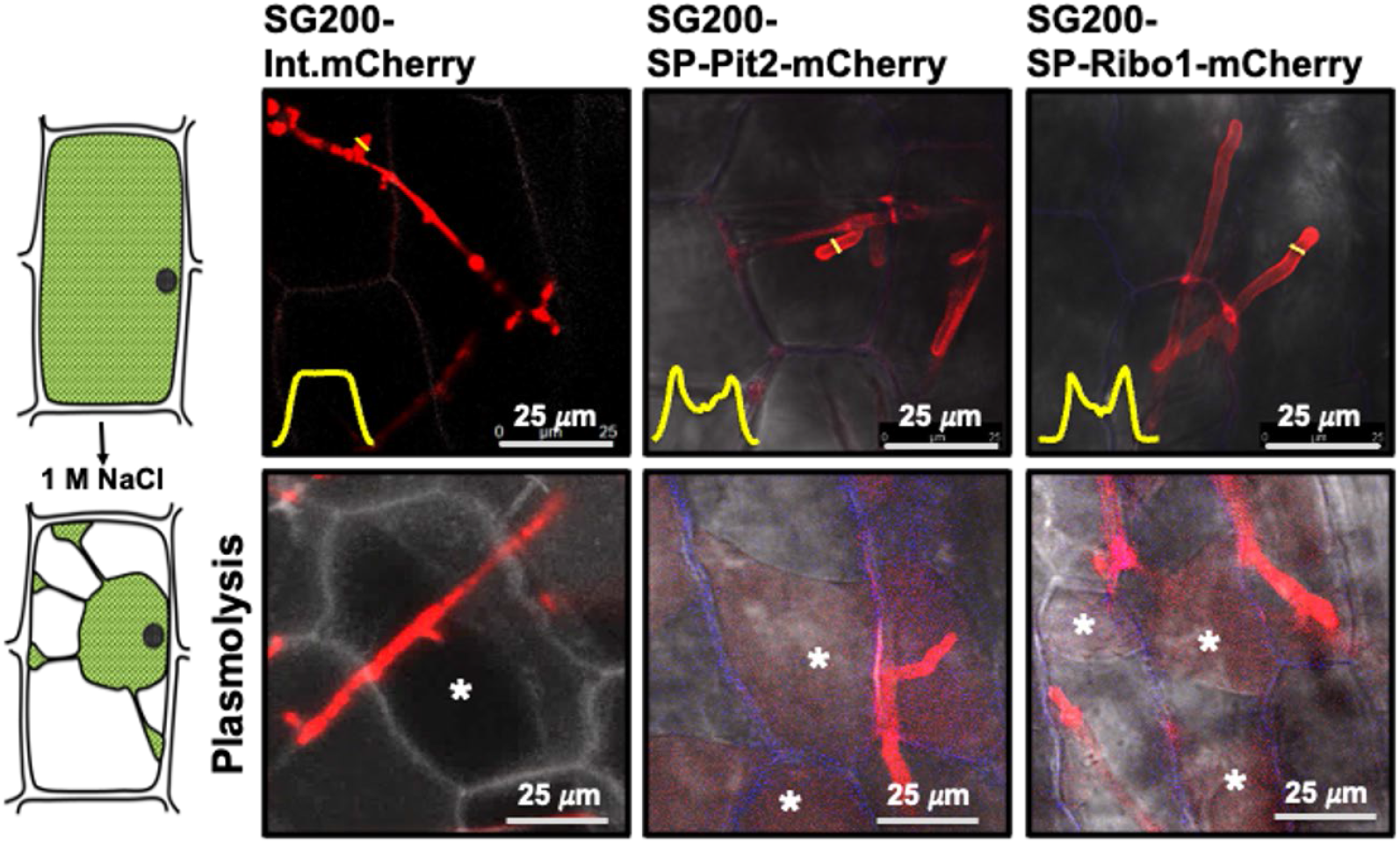
Localization of Ribo1-mCherry during host colonization. Ribo1-mCherry was heterologously expressed in *U. maydis* SG200 strain under control of the *pit2* promoter. The SG200 strains expressing the Ribo1-mCherry, UmPit2-mCherry (as a positive control for secretion) and cytosolic mCherry (int. mCherry; as a negative control for secretion) were inoculated on maize seedlings. Confocal microscopy was performed at 2 days post infection (dpi) to monitor the localization of each recombinant protein. The yellow graphs indicate the mCherry signal intensity along the diameter of the hyphae (illustrated by yellow lines in the image). Plasmolysis was performed with 1 M NaCl solution. White asterisks indicate apoplastic space after plasmolysis.

### Overexpression of Ribo1 negatively affects *Ustilago maydis* infection

To test whether Ribo1 induces host plant cell death in maize, we generated a *U. maydis* strain overexpressing both native and enzymatic inactive versions of Ribo1. To this end, both Ribo1 and UhRibo1-M^E80Q/D98N^ (hereafter referred to as Ribo1^M^) were overexpressed in *U. maydis* SG200 under control of *Umpit2* promoter. The overexpression of *Ribo1* genes was confirmed by RT-qPCR **(Fig. S2a)** and subsequently plant infection assays were performed with *U. maydis* overexpressing (OE) Ribo1 and Ribo1^M^ mutant strains **(Fig. S2b-c)**. In addition to single insertion of the OE constructs, *U. maydis* strains carrying multiple copies of the constructs were also used to test for a potential dose-effect of overexpressed Ribo1 during plant infection **(Fig. S2b)**. The plant infection assays revealed that overexpression of either Ribo1 or Ribo1^M^ hampers *U. maydis* colonization in maize **(Fig. 5a-b** and **Fig. S2b-c)**. While SG200-OE-Ribo1^M^ showed reduced virulence compared to SG200 progenitor strain; overexpression of the enzymatic active Ribo1 (strain SG200-OE-Ribo1) led to a severe reduction in virulence, suggesting an additional cytotoxic activity of the native protein **(Fig. S2b-c)**. *U. maydis* containing multiple integrations of *Ribo1* and *Ribo1*^*M*^ not only failed to form tumors (only few small tumors were observed in SG200-OE-Ribo1^M^-infected plants), but they also showed more necrotic spots on maize leaves, which indicates plant cell death **(Fig. 5b** and **Fig. S2c-d)**. This shows that Ribo1 has a dominant negative effect on *U. maydis* virulence in a dose-dependent manner. Moreover, infection assays performed with SG200-OE-Ribo1 (multiple integration) on 19 different maize lines showed a consistent avirulence phenotype with no tumor formation and formation of chlorotic and necrotic spots **(Fig. S2d)**. Southern blot analysis was performed to confirm the number of gene integration **(Fig. S2e)**. To test whether *U. maydis* overexpressing Ribo1 induces a specific maize defense response, alexa fluor (AF)-488 conjugated wheat germ agglutinin (WGA) and propidium iodide (PI) staining (WGA-AF488/PI), 3,3’-diaminobenzidine (DAB) and aniline blue staining were performed with maize leaves infected by the strains SG200, SG200-OE-Ribo1 and SG200-OE-Ribo1^M^. WGA-AF488/PI staining revealed hampered fungal growth of SG200-OE-Ribo1 compared to the progenitor strain SG200 **(Fig. 5c)**. Moreover, SG200-OE-Ribo1 infected leaves showed increased PI signals, which indicates enhanced cell wall fortifications in response to SG200-OE-Ribo1 **(Fig. 5c)**. Similarly, both DAB staining and aniline blue staining showed elevated ROS and callose accumulation in SG200-OE-Ribo1 strain infected maize leaves, respectively **(Fig. 5d-f)**. The aniline blue staining also revealed that the SG200-OE-Ribo1^M^ strain induced more callose deposition compared to SG200-OE-Ribo1 strain **(Fig. 5e-f)**. RT-qPCR on the maize *pathogenesis-related* genes *ZmPR1* and *ZmPR5* showed a significant induction in response to SG200-OE-Ribo1 compared to SG200 **(Fig. 5g)**. Maize leaves infected by SG200-OE-Ribo1^M^ also showed induced *ZmPR5* gene expression, however, for *ZmPR1* the observed induction appeared to be not significant **(Fig. 5g)**.

**Fig. 5.**
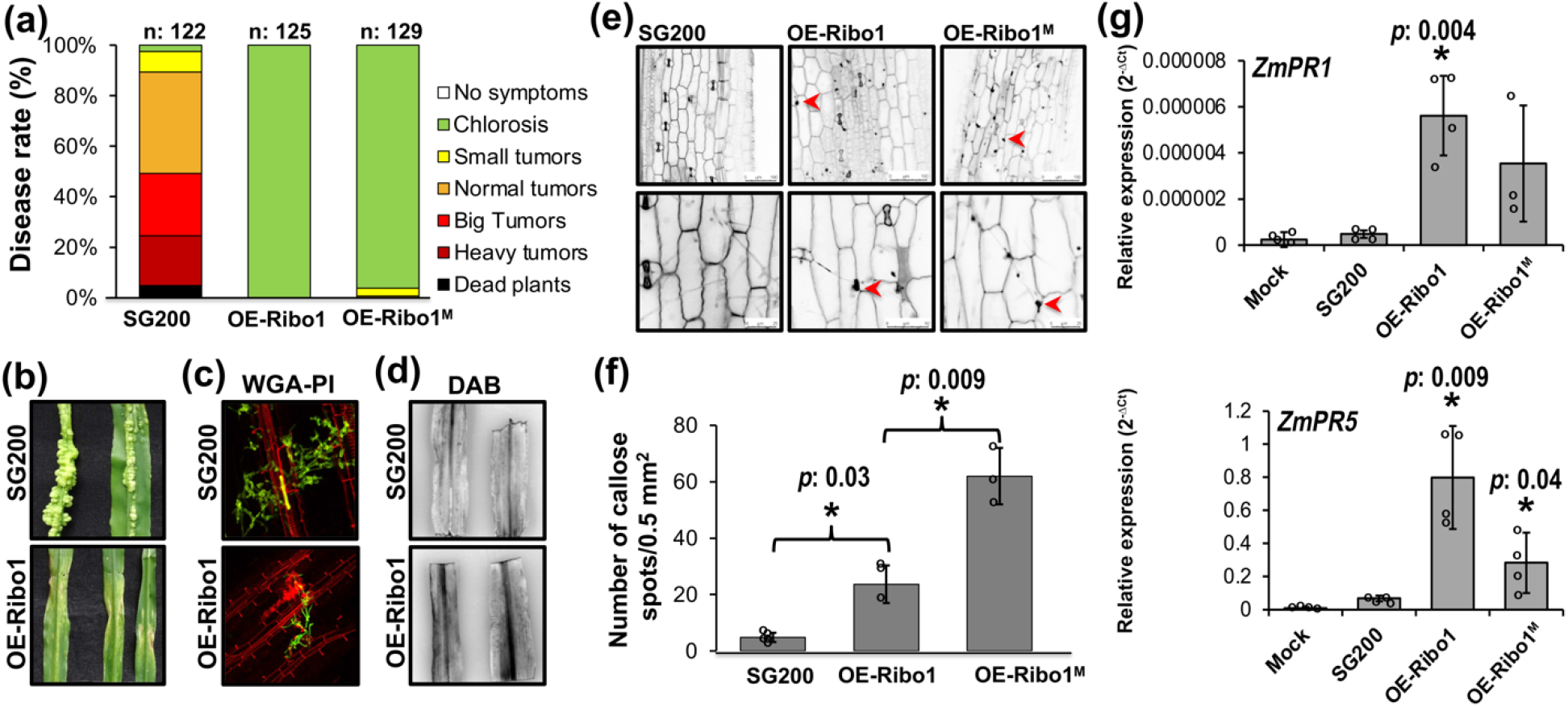
Overexpression of Ribo1 in *Ustilago maydis*. **(a-b)** Disease symptoms caused by *U. maydis* SG200, OE-Ribo1 (multiple integration) and OE-Ribo1^M^ (multiple integration) strains on Early Golden Bantam (EGB) maize leaves at 12 days post inoculation (dpi). Disease rates are given as a percentage of the total number of infected plants. Three biological replicates were carried out. n: number of infected maize seedlings. **(c)** Microscopic observation of *U. maydis* SG200 and OE-Ribo1 hyphae during maize colonization at 4 dpi via WGA-AF488/PI staining. WGA-AF488 (green color -fungal cell wall): excitation at 488 nm and detection at 500-540 nm. PI (red color - plant cell wall): excitation at 561 nm and detection at 580–630 nm. **(d)** 3,3’-diaminobenzidine (DAB) staining of *U. maydis* SG200 and OE-Ribo1 infected maize leaves. **(e-f)** Aniline blue staining for quantification of callose deposition in *U. maydis* SG200, OE-Ribo1 and OE-Ribo1^M^ infected maize leaves. Red arrows points callose depositions. **(f)** Quantification of callose deposition. At least three independent biological replicates were performed for callose deposition. Data were presented as mean value ± SD. Asterisks above bars indicate significant differences (two-tailed student’s t-test). *p*-values are illustrated on the graph. **(g)** RT-qPCR for *pathogenesis related (PR)* gene expression on SG200, OE-Ribo1 or OE-Ribo1^M^ strains infected maize leaves at 4 days post infection (dpi). The expression levels of maize *PR* genes, including *PR1* and *PR5* were calculated relative to the *GAPDH* gene of maize. Four independent biological replicates were performed for RT-qPCR. Data were presented as mean value ± SD. Asterisks above bars indicate significant differences (two-tailed student’s t-test). *p*-values are illustrated on the graph.

### Smut Ribo1 has antimicrobial activity

Analysis of publicly available transcriptome data of *U. maydis* and *U. hordei* showed that these smut fungi express Ribo1 only at very early stages of infection (12 hpi – 2 dpi) and that Ribo1 expression is strictly down regulated afterwards (Lanver *et al*., 2018; Ökmen *et al*., 2018b; Zuo *et al*., 2021) **(Fig. S3a-b)**. Its expression pattern together with the high cytotoxic activity led us to hypothesize that Ribo1 antagonizes microbial growth, rather than interfering with the plant host.

To test this idea, we produced Ribo1 using *Escherichia coli* and *Pichia pastoris* as protein expression systems without and with signal peptide, respectively. However, it appeared that both *E. coli* and *P. pastoris* strains expressing Ribo1 were significantly attenuated in growth, reflecting the cytotoxic activity of the protein **(Fig. 6a-c)**. Because of this antimicrobial activity, neither *E. coli* nor the *P. pastoris* protein expression system could produce recombinant Ribo1 protein. To ascertain if Ribo1 is engaged in competition with other host microbes, two bacteria were isolated from the maize leaf surface and their identity was determined by 16S rRNA sequencing. Ultimately, two bacterial species, *Pantoea dispersa* (Pd) and *Pantoea agglomerans* (Pa), were identified on the maize leaf surface. Co-inoculation of *U. maydis* SG200 with *Pd* and *Pa* mixture resulted in a reduction in tumor formation, indicating a negative effect of these two bacteria on fungal virulence **(Fig. S3c)**. Strikingly, although the SG200Δribo1 mutant did not show any significant difference in virulence compared to the SG200 **(Fig. S3d)**, in the disease assay performed with co-inoculation of the *Pd*-*Pa* mixture with the SG200 or the SG200Δribo1 mutant strains, the deletion mutant showed significant reduction in virulence compared to SG200 **(Fig. 6d)**. In co-culture *in vitro* experiments, constitutive expression of *Ribo1* under the control of *actin* promoter in *U. hordei* strain DS200 resulted in a significant growth inhibition of *P. dispersa* **(Fig. 6e)**. Together, we conclude that Ribo1 enables smut fungi to compete with other host associated microbes and thereby colonize their host niche.

**Fig. 6.**
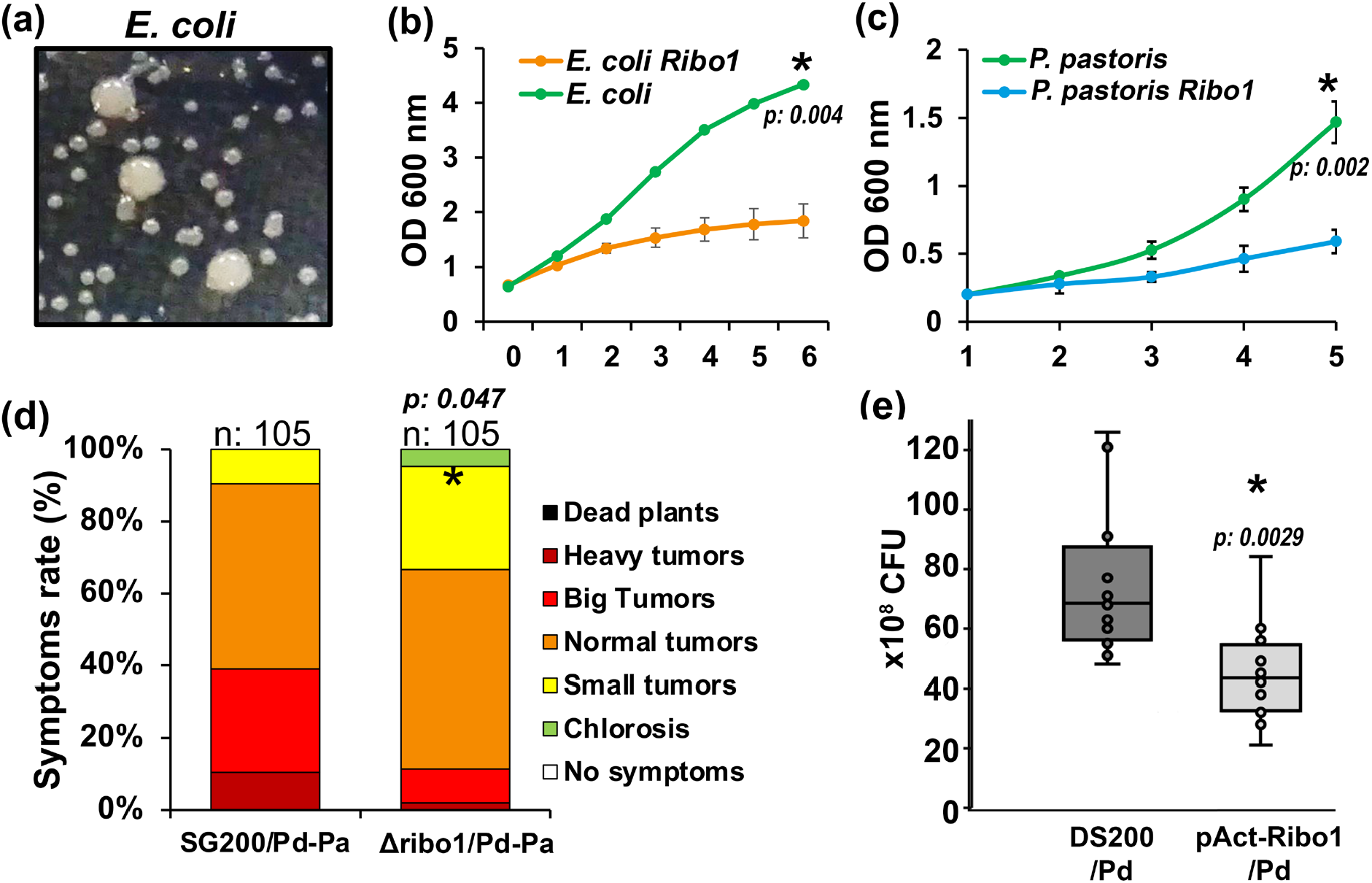
Ribo1 has antimicrobial activity. **(a)** *Escherichia coli* expressing Ribo1 protein has slower growth rate compared to empty vector control. While small colonies in the picture represent *E. coli* carrying Ribo1 integrated plasmid, bigger colonies represent empty plasmid. **(b)** Expression of Ribo1 in *Escherichia coli*. Growth rates of both *E. coli* with empty plasmid and *E. coli* with *Ribo1* expressing plasmid were monitored after IPTG induction. **(c)** Expression of Ribo1 in *Pichia pastoris*. Growth rates of both *P. pastoris* and *P. pastoris* expressing *Ribo1* were monitored. Data were presented as mean value ± SD. Asterisks above bars indicate significant differences (two-tailed student’s t-test). *p*-values are indicated on the graph. **(d)** *In vitro* growth inhibition assay for *Pantoea. U. hordei* DS200 or DS200-pAct::Ribo1 (OD: 1) (for constitutive gene expression) strains were co-incubated with *P. dispersa* (OD: 0.2) in YEPS_Light_ medium for four hours and subsequently plated on LBA medium. To ensure that only *Pantoea* colonies were counted, cycloheximide (an eukaryotic-specific protein synthesis inhibitor) was added to the LBA medium, thereby inhibiting DS200 colony formation. Results were recorded as colony forming units (CFUs) of Pantoea and three biological replicates were carried out. Three independent biological replicates were performed for growth inhibition assay. Statistical significance was determined using a two-tailed student’s t-test, with *p*-values indicated on the bars and asterisks above the bars representing significant differences. **(e)** Impact of maize associated *Pantoea* sp. on *Ustilago maydis* colonization. Disease symptoms caused by SG200 and SG200Δribo1 mutant strains mixed with *P. dispersa* (*Pd*) and *P. agglomerans* (*Pa*) on Early Golden Bantam (EGB) maize leaves at 10 days post inoculation (dpi). Disease rates are given as a percentage of the total number of infected plants. Three biological replicates were carried out. n: number of infected maize seedlings. Asterisks above bars indicate significant differences (two-tailed student’s t-test). *p*-values are indicated on the bars.

## Discussion

Plant pathogenic fungi rely on an arsenal of virulence factors (so called effectors), which are expressed and delivered into the host tissue to enable colonization of the plant and fungal proliferation (Stergiopoulos & de Wit, 2009; Lo Presti *et al*., 2015). It has been shown that the effector repertoire of a pathogen also determines its pathogenic life style. While necrotrophs possess relatively more plant cell wall degrading enzymes and cell death inducing toxins to kill their respective host cells, biotrophs have effector set to suppress plant cell death since they are strictly relying on live host cells (Zhao *et al*., 2013; Rodriguez-Moreno *et al*., 2018). Like other biotrophs, smut fungi also have restricted number of plant cell wall degrading enzyme to keep their respective host cell intact and alive; however, in this study we have identified and functionally characterized a conserved, small secreted protein from smut fungi that has a ribonuclease T1 (RNase T1) domain and broad-spectrum cytotoxic activity.

### A conserved secreted ribonuclease from smut fungi induces plant cell death

Extracellular ribotoxins specifically cleave conserved sarcin/ricin loop of rRNA leading to inhibition of protein biosynthesis and cell death (Garcia-Ortega *et al*., 2002; García-Mayoral *et al*., 2005; Citores *et al*., 2018). Although our phylogenetic tree analysis showed that smut Ribo1 is closer to non-toxic RNase T1 members, heterologous expression of the *U. hordei, U. maydis*, or *S. reilianum* Ribo1 with or without secretion signal had cell death-inducing activity in *N. benthamiana*. Moreover, Ribo1 of the hemibiotrophic maize pathogen *F. verticillioides* also showed plant cell death-inducing activity on tobacco, indicating a conserved function of Ribo1 among these phytopathogenic fungi. Overexpression of *Ribo1* in *U. maydis* SG200 also caused cell death induction on infected maize plant indicating cytotoxic activity of Ribo1 on both host and non-host plants. However, the mode of action of cell death-inducing activity of secreted smut Ribo1 is still elusive.

### Extracellular Ribo1 induces plant cell death independent from enzymatic activity

It is well reported that conserved Nep1-like proteins (NLPs) from a wide range of fungal and oomycete phytopathogens act as both toxin-like virulence factors and as elicitors of plant defense responses (Qutob *et al*., 2006; Ottmann *et al*., 2009; Böhm *et al*., 2014; Lenarčič *et al*., 2017), which is consistent with our findings of Ribo1 on maize and tobacco plants. Furthermore, the RALPH effectors of *B. graminis* f. sp. *hordei*, are recognized by barley (Praz *et al*., 2017), although they were outliers in the constructed phylogenic tree, their structural similarity to Ribo1 may indicate that also Ribo1 could act as MAMPs to induce plant immunity. One can also hypothesize that the Ribo1-induced cell death in *N. benthamiana* could result from cleavage of tobacco rRNA. However, while both secreted and intracellularly expressed Ribo1 induce plant cell death, only the secreted version of UhRibo1-M^E80Q/D98N^ does so. This suggests that Ribo1-induced cell death inside the plant cell is associated with its enzymatic activity, while the secreted Ribo1 is likely acting as an elicitor of plant cell death. Kettle *et al*., (2018) reported that the Zt6 ribonuclease from *Zymoseptoria tritici*, the causal agent of Septoria leaf blotch disease of wheat, induces cell death in tobacco independently of Brassinosteroid Insensitive 1 (BRI1)-Associated Receptor Kinase 1 (BAK1) and Suppressor of BIR1-1 (SOBIR1) (Kettles, Graeme J. *et al*., 2018). Consistently, we found that the cell death induced by smut Ribo1 is also SOBIR1 independent.

### N-terminal 11 amino acids are essential for Ribo1 cytotoxicity

Previously, it has been shown that the N-terminal region of secreted ribotoxins is important for their cellular toxicity (Garcia-Ortega *et al*., 2002; García-Mayoral *et al*., 2005). Compared to non-toxic RNase T1, ribotoxins possess longer N-terminal loops in a β-sheet–hinge-β-sheet structure that plays an important role in ribosome binding and cellular uptake, which is essential for their cytotoxicity (Garcia-Ortega *et al*., 2002; García-Mayoral *et al*., 2005). For example, deletion of the N-terminal 7-22 amino acids from *Aspergillus* sp. alpha-sarcin results in less efficient interaction with both the ribosome and cell membrane, impairing cellular uptake (Garcia-Ortega *et al*., 2002). In this study, we found that a splice variant of FvRibo1, which lacks 24 N-terminal amino acids (from 38 to 61th amino acid), was unable to induce cell death. Accordingly, deletion of a conserved 11 amino acid motif in *U. hordei* Ribo1 also resulted in loss of cell death induction *in planta*, indicating importance of this region for cytotoxicity. 3-D structural prediction of FvRibo1_L and FvRibo1_S revealed that the N-terminal β-sheet–hinge-β-sheet part was completely abolished in FvRibo1_S spliced version, highlighting the importance of this part for cytotoxicity **(Fig. 3b)**.

### Ribo1 induces immune responses in smut hosts

The primary function of effectors secreted by biotrophic phytopathogens is to inhibit host defense responses and particularly programmed cell death, which restricts the fungal growth and host colonization (Lo Presti *et al*., 2015). Therefore, the presence of a plant cell death inducing secreted protein in the biotrophic smut fungi appears to be surprising. However, transcriptome data of different smut fungi shows that Ribo1 is only expressed at early time points (at 1 and 2 dpi), which coincide with fungal differentiation on the plant surface. Upon host penetration and during all stages of *in-planta* growth, expression of *Ribo1* is strictly downregulated (Lanver *et al*., 2018; Ökmen *et al*., 2018b; Zuo *et al*., 2021). Accordingly, the overexpression of Ribo1 driven by the promoter of the Pit2 effector, an essential virulence factor, drastically reduced fungal colonization in a dose-dependent manner. Both multiple integration strains of SG200-OE-Ribo1 and SG200-OE-Ribo1^M^ lost the ability to induce tumor formation and triggered the formation of necrotic lesions on infected maize leaves. Significant reduction of virulence in SG200-OE-Ribo1^M^ strain expressing inactive Ribo1 also suggests that not only the cytotoxic activity of the Ribo1 negatively affects the fungal colonization, but also that recognition of Ribo1 induces plant defense responses.

Consistent with this hypothesis, SG200 overexpressing either Ribo1 or Ribo1^M^ induced several defense related responses including cell wall fortification, ROS accumulation, callose deposition and *PR* gene expression on maize plant. Furthermore, SG200-OE-Ribo1^M^ strain showed higher callose deposition compared to both SG200 and SG200-OE-Ribo1 strains suggesting that the cytotoxic activity of the Ribo1 protein may lead to a reduced induction of defense responses. Consistent with these data, we recently reported that overexpression of *F. verticillioides* Ribo1 in *U. hordei* negatively affects its colonization and virulence during barley colonization (Ökmen *et al*., 2021). Similarly, expression of *Zymoseptoria tritici* Zt6 sRNase in wheat plants induces cell death, which is in line with necrotic spots formation in SG200-OE-Ribo1 infected maize leaves (Kettles, G. J. *et al*., 2018).

### Ribo1 engages in competition with the host microbial community

Knock out of *Ribo1* in *U. maydis* did not result in a significant reduction in fungal virulence, indicating that it does not play a major role during *U. maydis* infection and tumorigenesis **(Fig. S3d)**. Deletion of the ribonuclease *Zt6* in *Z. tritici* also did not lead to any impaired virulence phenotype on susceptible wheat plant (Kettles, G. J. *et al*., 2018). However, the question still remains about the biological function of Ribo1 in plant colonizing fungi. Our multiple attempts to produce recombinant Ribo1 protein in *E. coli* or in *P. pastoris* revealed that the protein exhibits cytotoxic activity against both bacteria and yeast. In line with this, ribotoxins from *Aspergillus* sp. and from *Z. tritici* ware also found to have a strong cytotoxic activity against bacterial, fungal and insect cells (Citores *et al*., 2018; Kettles, Graeme J. *et al*., 2018). Moreover, the fact that transcriptional activation of *Ribo1* in smut fungi is specifically restricted to the very early stage of infection when the fungus grows on the plant surface, suggests a role in competition with other host-associated microbes rather than with the host plant.

To follow this up, we isolated bacteria from maize leaves, which led to the identification of two dominating species, *P. dispersa* (*Pd*) and *P. agglomerans* (*Pa*). *Pantoea* sp. has previously been reported to be an effective biocontrol agent against black rot in sweet potato (Jiang *et al*., 2019). In accordance with these findings, we found that co-inoculation of a mixture of *Pd*-*Pa* with *U. maydis* SG200 strain negatively affected tumor formation, which is the indication of successful fungal colonization. Moreover, we found that the *U. maydis* Δribo1 mutant was even more strongly affected than the wildtype strain by co-inoculation with a *Pd*-*Pa* mixture. This reduced fitness of the deletion strain in presence of the leaf-associated bacteria suggests that Ribo1 contributes to the ability of smut fungi to compete with other plant-associated microbes and thereby occupy their own niche in the host. Furthermore, we also found that constitutive expression of Ribo1 in smut fungi restricts growth of *P. dispersa in vitro*, which confirms the antimicrobial activity of this protein.

The finding that smuts secrete a cytotoxic Ribo1 contributing to the pathogen’s fitness in competition with other host-associated microbes is consistent with recent findings made for the wilt pathogen *Verticillium dahlia*, which deploys effector proteins that modulate the host microbiota (Snelders *et al*., 2018; Snelders, NC *et al*., 2021). Taken together, our observations emphasize the emerging role of fungal effector proteins in the modulation of the microbial communities, which increases the pathogen’s biological fitness within the plant holobiont.

## Materials and Methods

### Fungal and plant growth conditions

The *Escherichia coli* DH5α strain was grown in dYT-liquid medium (1.6% w/v peptone, 1% w/v yeast extract and 0.5% w/v NaCl) with appropriate antibiotics at 37°C with 200 rpm shaking. *Ustilago maydis* SG200 solopathogenic strain was incubated in YEPS_light_ (0.4% w/v yeast extract, 0.4% w/v peptone, and 2% w/v sucrose) liquid medium at 28°C and 20°C with 200 rpm shaking, respectively. Growth of *U. maydis* culture on plates was carried out on potato dextrose agar with appropriate antibiotics (concentration: 2 µg/ml carboxin). The *P. dispersa* and *P. agglomerans* were grown in dYT-liquid medium (1.6% w/v peptone, 1% w/v yeast extract and 0.5% w/v NaCl) at 28°C with 200 rpm shaking. The *Pichia pastoris* KM71H-OCH strain was used for recombinant protein expression. YPD liquid medium supplemented with 100 µg/ml zeocin was used for the initial growth of *P. pastoris* strains at 28°C and 200 rpm shaking. *Zea mays* L. Early Golden Bantam (EGB) maize (Olds Seeds, Madison, WI, USA) and *Hordeum vulgare* Golden Promise (GP) barley cultivars were used for infection assays. GP cultivar was grown in a greenhouse at 70% relative humidity, at 22°C during the day and the night; with a light/dark regime of 15/9 hrs and 100 Watt/m^2^ supplemental light when the sunlight influx intensity was less than 150 Watt/m^2^.

### Nucleic acids methods

All plasmid DNA isolation from bacterial cells was performed using the QIAprep Spin Miniprep Kit (Qiagen; Hilden, Germany) according to the manufacturer’s information. The *U. maydis* genomic DNA isolation was performed according to the protocol described by Schultz *et al*. (1990). Phusion© polymerase enzyme (Thermo Fisher Scientific;Darmstadt, Germany) was used in the polymerase chain reaction (PCR) to amplify specific DNA fragments by using gene specific primer pairs depicted in **Supporting Information File 1**.

Total RNA was isolated from crushed infected leaf material (at 4 dpi) using the TRIzol® extraction method (Invitrogen; Karlsruhe, Germany) according to the manufacturer’s instructions. Genomic DNA contamination from total RNA was removed by using the Turbo DNA-Free™ Kit (Ambion/Applied Biosystems; Darmstadt, Germany). cDNA synthesis was performed by using the First strand cDNA synthesis Kit with 1 µg of total isolated RNA (Thermo Fisher Scientific; Darmstadt, Germany). RT-qPCR analysis for *PR* gene expression was performed by using SYBR® Green Supermix (BioRad; Munich, Germany). The following program was used for the RT-qPCR reaction in a Bio-Rad iCycler system: 2 min at 95°C followed by 45 cycles of 30 s at 95°C, 30 s at 61°C and 30 s 72°C. The expression levels of maize *PR* genes were calculated relative to maize *GAPDH* gene (*NM001111943*). Results of at least three biological RT-qPCR replicates were analyzed using the 2^−ΔCt^ method (Livak & Schmittgen, 2001). The primers used for RT-qPCR are depicted in **Supporting Information File 1**.

### Construction of over expression vectors

For heterologous gene expression constructs (*p123-pUmPit2::SP-Ribo1, p123-pUmPit2::SP-Ribo1*^*M*^ and *p123-pUmPit2::SP-Ribo1-mCherry*), standard molecular biology methods were used according to molecular cloning laboratory manual of Sambrook *et al*. (1989) (Sambrook *et al*., 1989). Amplified PCR fragments for each gene of interest were digested with appropriate restriction enzymes and subsequently ligated into selected destination vector that was also digested with the same restriction enzymes by using T4-DNA ligase (New England Biolabs; Frankfurt a.M., Germany) according to manufacturer’s instructions. All primer pairs and restriction sites for cloning of each vector are listed in **Supporting Information File 1**. *E. coli* transformation was performed via heat shock assay according to standard molecular biology methods (Sambrook *et al*., 1989). The sequences confirmation of each construct was performed via sequencing at the Eurofins Genomics (Cologne, Germany). Golden gate cloning system was used to clone Ribo1 in to binary vector of *pL1M* (Engler *et al*., 2014). *UHOR_02675* active site mutants (Ribo1-M^E80Q/D98N^) were created from by using QuickChange XL Site-Directed Mutagenesis Kit from Agilent Technology according to manufactures instructions. All primer pairs and restriction sites for cloning of each vector are listed in **Supporting Information File 1**.

### *Agrobacterium*-mediated transient expression of Ribo1 in tobacco

All selected *Ribo1* homologs from different fungal species were cloned into the *p1LM* golden gate expression vector under the control of *35SCaMV* promoter and with C-terminally HA-tag. *Agrobacterium*-mediated transient transformation assay (ATTA) was performed by using *Agrobacterium tumefaciens* GV3101 strain on wildtype and Δsobir1 *N. benthamiana* leaves. ATTA assay was performed according to van der Hoorn *et al*., 2000 (Van der Hoorn *et al*., 2000). *Agrobacterium* strains carrying different *Ribo1* gene were infiltrated into 5-6 week-old *N. benthamiana* leaves with a syringe without needle. *Agrobacterium* strain carrying *eGFP* was used as a negative control for cell death induction. Ribo1-mediated cell death induction was observed at 4-5 dpi. The Δsobir1 *N. benthamiana* seeds were kindly provided by Dr. M.H.A.J. Joosten from Laboratory of Phytopathology, University of Wageningen.

### CRISPR/Cas9 gene editing system

The CRISPR/Cas9-HF (high fidelity) gene editing system was used to knockout *U. maydis* Ribo1 according to Zuo et al. (2020) (Zuo *et al*., 2020). The sgRNA for the knockout of the *U. maydis Ribo1* gene was designed by e-CRISPR (http://www.e-crisp.org/ECRISP/aboutpage.html) **(Supporting Information File 1)** (Heigwer *et al*., 2014). The sgRNA for *Ribo1* gene was expressed under the *pU6* promotor. Linearized CRISPR/Cas9-HF plasmid (with Acc65I restriction enzyme) was assembled with spacer oligo and scaffold RNA fragment with 3’ downstream 20 bp overlap to the plasmid using Gibson Assembly (Gibson *et al*., 2009).

### Fungal transformation and disease assay

The *U. maydis* transformation assay was performed by using protoplasts according to Kämper, 2004. Disease assays for SG200, SG200Δribo1, SG200-OE-Ribo1 and SG200-OE-Ribo1^M^ strains were performed according to Ökmen *et al*., 2018a. Briefly, all *U. maydis* strains were grown in YEPS_light_ liquid medium at 28°C and 200 rpm shaking until getting an OD_600_ of 0.6-0.8. Subsequently, *U. maydis* cells were centrifuged at 3000 rpm for 10 min at RT and resuspended in sterile distilled water to an OD_600_ of 1.0. Then each *U. maydis* cell suspension was injected into stems of 7-day-old maize seedlings (Early Golden Bantam) with a syringe with needle. All infection assays were performed at least in three biological replicates. To confirm that *U. maydis* can overexpress Ribo1 gene, RT-qPCR was performed with SG200, SG200-OE-Ribo1 and SG200-OE-Ribo1^M^ strains infected maize leaves. All primer pairs are listed in **Supporting Information File 1**. To apply statistical analysis on *U. maydis* disease assays, the disease index was calculated as follows: The number of plants sorted into categories ‘chlorosis’, ‘small tumor’, ‘normal tumor’, ‘big tumor’ and ‘heavy tumor’ were multiplied by 1, 3, 6, 9 and 12, respectively. All calculated numbers for each strain were summed and then divided by the total number of infected plants to obtain the disease index.

### Localization of Ribo1-mCherry during host colonization

To visualize secretion of Ribo1-mCherry in *U. maydis* during maize colonization, Ribo1-mCherry expressing *U. maydis* strain was inoculated on maize leaves. Subsequently, infected maize leaves were monitored by using a Leica SP8 confocal microscopy for localization of Ribo1-mCherry at 3 dpi. Pit2-mCheery and Int. mCherry expressing *U. maydis* strains were used as a positive and negative controls for extracellular secretion, respectively. An excitation at 561 nm and detection at 580–630 nm was used to detect mCherry signals.

### WGA-AF488/Propidium iodide staining

*U. maydis* infected leaves were stained with the wheat germ agglutinin (WGA)-AF488 (Molecular Probes, Karlsruhe, Germany) and propidium iodide (Sigma-Aldrich) according to Ökmen *et al*. (2018b). While WGA-AF488 stains fungal chitin cell walls (green), the propidium iodide stains host plant cell walls (red). After infected leaf materials were bleached in pure ethanol, they were boiled for 2-3 hours in 10% KOH at 85°C. Subsequently, the pH of samples was neutralized using 1xPBS buffer (pH: 7.4) with several washing steps. The WGA-AF488/PI staining solution (1 µg/ml propidium iodide, 10 µg/ml WGA-AF 488; 0.02% Tween 20 in PBS pH 7.4) was vacuum infiltrated into infected leaf material for three times 5 min with a desiccator at 250 mbar. WGA-AF488: excitation at 488 nm; detection at 500-540 nm. PI: excitation at 561 nm; detection at 580– 630 nm.

### Aniline blue staining

After infected maize leaves were first fixed and decolorized in acetic acid: ethanol (v/v; 1:3) mixture, all samples were vacuum infiltrated with 150 mM K_2_HPO_4_ three times for 10 min. Subsequently, the buffer was replaced with 0.05% aniline blue solution (w/v in 150 mM K_2_HPO_4_) and the stain was vacuum infiltrated in the leaves three times for 10 min with a desiccator at 250 mbar. Aniline blue infiltrated infected maize leaves were incubated at room temperature at dark for 1 hour and the leaf samples were monitored by using confocal microscopy with DAPI channel (Emission 490-520 nm).

### The 3,3′-diaminobenzidine (DAB) staining

The 3,3′-diaminobenzidine (DAB) staining was performed according to a previous methods reported by Daudi and O’Brain 2012 (Daudi & O’Brien, 2012). SG200 and SG200-OE-Ribo1 strain infected maize leaves (at 4 dpi) were vacuum infiltrated with DAB/Mn2+ staining solution containing 0.2% DAB-tetrahydrochloride, 20 mM MnCl24H2O, 1 mM sodium azide and 0.05% Tween20. After incubation of 8 hours at dark, the DAB/Mn2+ staining solution was replaced with bleaching solution (ethanol:acetic acid:glycerol, v/v/v 3:1:1) and photographs were taken after distaining of chlorophylls.

### Recombinant protein expression and purification

The *Pichia pastoris* KM71H-OCH protein expression system was used to produce C-terminally His tagged Ribo1 and Ribo1^M^ recombinant proteins. *Ribo1* and Ribo1^M^ genes were cloned into pGAPZαA vector (Invitrogen; Carlsbad, USA) under the control of a constitutive promotor with an α-factor signal peptide for secretion. Recombinant protein expression was performed in 1 L buffered (100 mM sodium phosphate buffer, pH 6.0) YPD medium at 28°C for 48 hours with 200 rpm shaking according to manufacturer’s instructions (pGAPZαA, B, & C *Pichia pastoris* Expression Vectors, Invitrogen; Carlsbad, USA). Protein purification was performed by using Ni-NTA-matrix according to manufacturer’s instructions (Ni-Sepharose™ 6 Fast-Flow, GE-Healthcare; Freiburg, Germany). After protein purification, protein samples were applied to NAP-25 buffer exchange column equilibrated with 20 mM potassium phosphate buffer pH 6.0 according to manufacturer’s instructions. The Ribo1 and Ribo1^Act.Mut^ recombinant proteins were stored at -20°C for further experiments.

### Microbial toxicity assays

To test Ribo1 has negative effect on growth rate of both *E. coli* and *P. pastoris*, both Ribo1 expressing *E. coli* and *P. pastoris* culture were started. *E. coli* and *P. pastoris* containing empty contracts were used as negative control. The protein expression in *E. coli* was induced by addition of 0.5 mM IPTG at OD:0.6 and the OD of cultures were measured every hour. The Ribo1 expressing *P. pastoris* was grown in YPD medium supplemented with 2% glucose and after dilution of the culture to OD:0.2, the OD of cultures were measured every two hours. All the collected values were at the end plotted on graph for visual representation.

### Isolation of epiphytic *Pantoae sp*. from maize leaf and inhibition assays

The *P. dispersa* (*Pd*) and *P. agglomerans* (*Pa*) were isolated by shaking out 0.5 g maize leaves in 4 ml of water supplemented with 0.1% tween-20 for two hours. Collected wash solution was plated on dYT-agar medium and incubated at 28°C for 2-4 days. The identity of isolated bacteria was determined by sequencing of their 16S rRNA using universal 16S rRNA primer pair **(Supporting Information File 1)**. Colony PCR was performed with the respective primer pair and purified PCR fragments were sequenced at LGC-Genomics. Co-inoculation assay with *U. maydis* SG200 and SG200Δribo1 strains were performed by having final OD of 1 for smut fungus and OD of 0.3 for each bacterial species. Subsequently, each culture mixtures including SG200, SG200+Pd/Pa and SG200Δribo1+Pd/Pa cell suspensions were injected into stems of 7-day-old maize seedlings with a syringe with needle. All infection assays were performed in three biological replicates.

For *in vitro* inhibition assay, OD of 1 *U. hordei* DS200 or DS200-pAct::Ribo1 (DS200 expressing *Ribo1* under the control of actin promoter for constitutive gene expression) strains were co-incubated with OD of 0.2 *P. dispersa* in YEPS_Light_ medium for 4 hours at 22°C with shaking. Subsequently, different dilutions of these co-cultures were plated on LBA supplemented with 0.01% cycloheximide to inhibit DS200 colony formation, thus only *Pantoea* sp. colonies could be counted. *In vitro* inhibition assays were performed in three biological replicates.

### Phylogenetic analysis

The amino acid sequence of Ribo1 (UHOR_02675) was blasted against NCBI the database by using BLASTp to find Ribo1 homologs in different fungi and bacteria. To construct a phylogenetic tree, the amino acid sequences of all selected Ribo1 homologs were aligned by using ClustalOmega (Sievers *et al*., 2011) and edited in Genedoc software (Nicholas *et al*., 1997). Subsequently, a consensus phylogenetic tree was constructed by using the minimum evolution algorithm with default settings and 1000 bootstrap replications in Mega7 software (Tamura *et al*., 2011). The *B. graminis tritici* RALFs effectors were used as an outgroup.

## Acknowledgements

BÖ acknowledges support from the Deutsche Forschungsgemeinschaft (DFG, German Research Foundation) OE 745/2-1 Project ID: 460397201. GD acknowledges support from the Cluster of Excellence on Plant Sciences (CEPLAS) funded by the Deutsche Forschungsgemeinschaft (DFG, German Research Foundation) under Germany’s Excellence Strategy – EXC 2048/1 – Project ID: 390686111. The authors would also like to thank Dr. M.H.A.J. Joosten from Laboratory of Phytopathology, Wageningen University for providing the Δsobir1 *N. benthamiana* seeds. The authors also thank Dr. Libera Lo Presti from the Cluster of Excellence CMFI University of Tübingen for constructive comments on the manuscript. We also thank Prof. Dr. Eric Kemen (Department of Microbial Interactions, IMIT/ZMBP, University of Tübingen) for generous support of this work

## Author contributions

B.Ö. and G.D. conceived the project. P.K. carried out golden gate cloning of Ribo1 for tobacco expression; R.W. performed southern blot assay; B.Ö. performed transformation, disease assays, protein production and microscopy. B.Ö. wrote the manuscript with input from all authors.

## Supporting Information

**Supporting Information File 1. Table for all constructs and primers. Supporting Figures**

**Fig. S1** Heterologous expression of *Ribonuclease T1* (*Ribo1*) in Δsobir1 *Nicotiana benthamiana*.

**Fig. S2** Maize leaves after infection with *U. maydis* strain overexpressing Ribo1.

**Fig. S3** Expression pattern of *Ribo1* genes in smut fungi.

