## Supporting Figures for "A conserved extracellular ribonuclease with broad-spectrum cytotoxic activity enables smut fungi to compete with host-associated bacteria"

**ORCIDs:**

Bilal Ökmen: https://orcid.org/0000-0002-4729-9973

Gunther Doehlemann: https://orcid.org/0000-0002-7353-8456

**Key words:** Effectors, ribotoxin, antimicrobial protein, smut fungi, cell-death induction

**Supporting Information File 1. Table for all constructs and primers.**

**Supporting Figures**

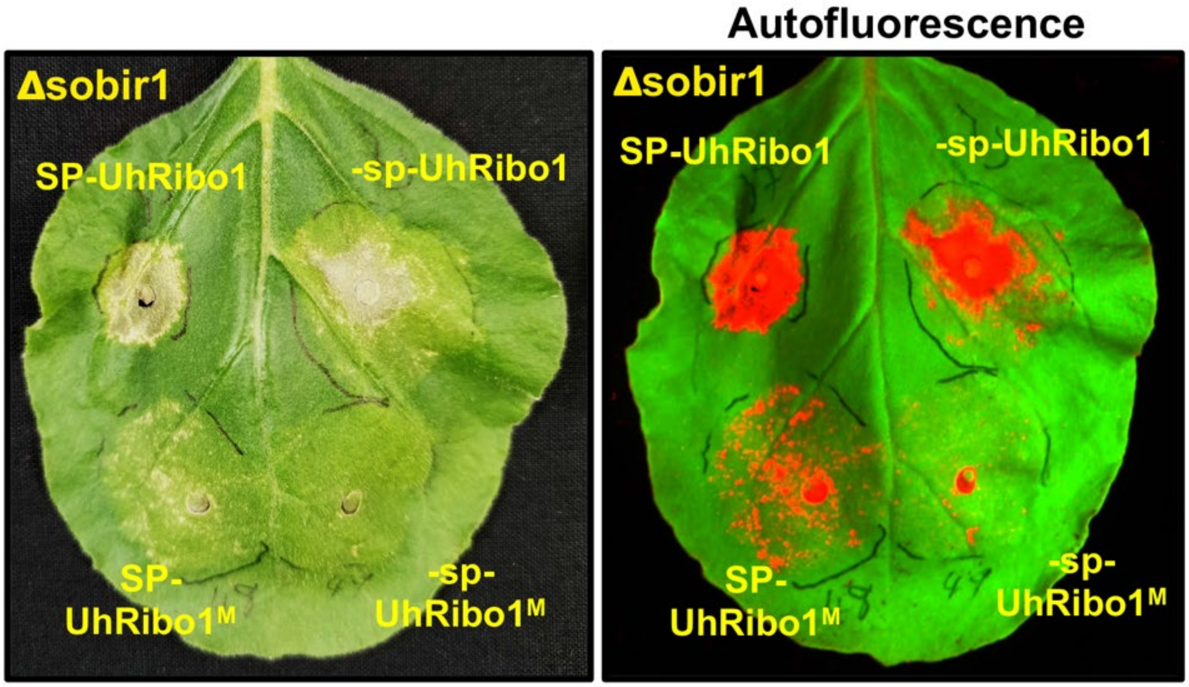


**Fig. S1 Heterologous expression of *Ribonuclease T1* (*Ribo1*) in Δsobir1 *Nicotiana benthamiana*.** By using the *A. tumefaciens*-mediated transient expression assay, the *UhRibo1* and active site mutant of *UhRibo1* (UhRibo1$^{M}$) were expressed in Δ*sobir1* tobacco plants with or without plant signal peptide (±SP). Picture were taken after 5 dpi. Autofluorescence pictures were taken with Bio-Rad ChemiDoc imaging system.

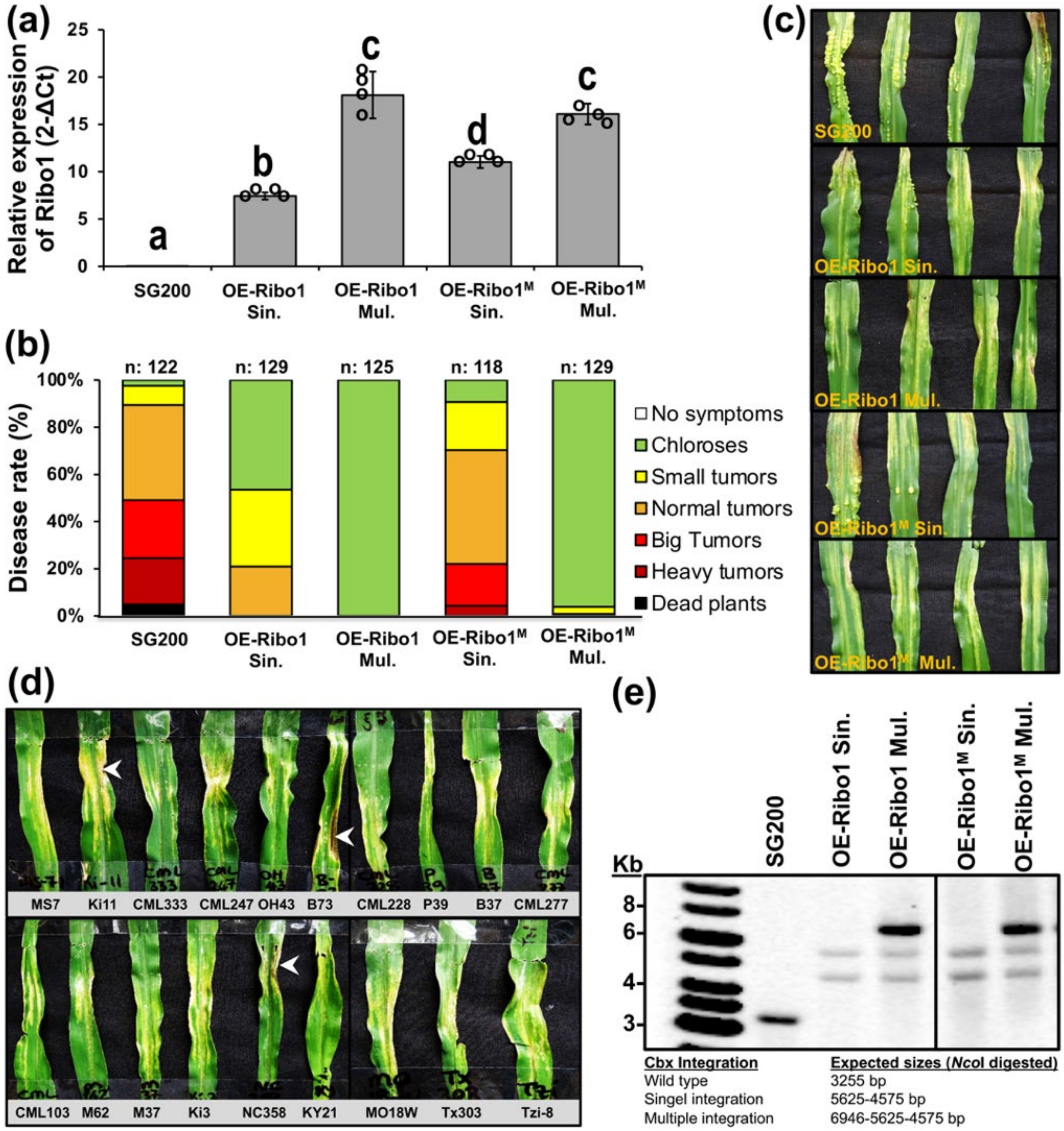


**Fig. S2 Maize leaves after infection with *U. maydis* strain overexpressing Ribo1. (a)** Relative expression of Ribo1 in SG200 and SG200 OE-Ribo1 mutant strains was measured by using RT-qPCR. M: active site mutant, Sin.: single gene integration, Mul.: Multiple gene integration. Four independent biological replicates were performed for RT-qPCR. Data were presented as mean value ± SD. Letters above bars indicate significant differences (two-tailed student's t-test). *UMAG_Ppi* gene was used as reference gene. **(b-c)** Disease symptoms caused by *Ustilago maydis* SG200, OE-Ribo1-Sin., OE-Ribo1-Mul., OE-Ribo1$^{M}$-Sin. and OE-Ribo1$^{M}$-Mul. strains on Early Golden Bantam (EGB) maize leaves at 12 days post inoculation (dpi). Disease rates are given as a percentage of the total number of infected plants. Three biological replicates were carried out. n: number of infected maize seedlings. OE of *Ribo1* in

SG200 hampers the colonization of the fungus in dose-dependent manner. M: active site mutant, Sin.: single gene integration, Mul.: Multiple gene integration. **(d)** Disease assay performed with 19 different maize cultivars infected by SG200 OE-Ribo1-Mul. White arrow head shows necrotic spots on maize leaves. **(e)** Southern blot analysis was performed to confirm SG200-OE-Ribo1 and SG200-OE-Ribo1$^{M}$ complementation. gDNA of both SG200 and complementation strains were digested with appropriate restriction enzyme. *Cbx* gene was used as a probe to detect single insertion event. Expected sizes for each insertion event depicted below figure.

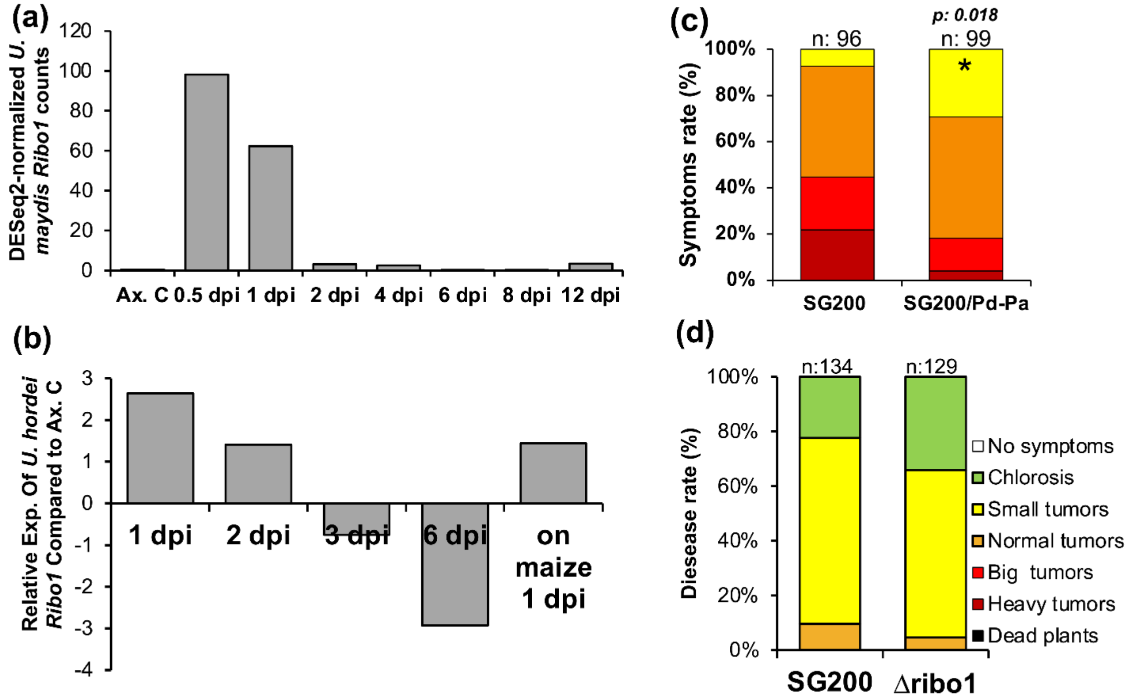


**Fig. S3 Expression pattern of *Ribo1* genes in smut fungi**. The bar graphs were created by using transcriptome data for **(a)** *U. maydis Ribo1* (Lanver *et al.*, 2018) and **(b)** *U. hordei Ribo1* (Ökmen *et al.*, 2018). **(c)** Impact of maize associated *Pantoea* sp. on *Ustilago maydis* colonization. Disease symptoms caused by *U. maydis* SG200 and SG200 (OD: 1) mixed with *Pantoea dispersa* (*Pd*) (OD: 0.3) and *Pantoea agglomerans* (*Pa*) (OD: 0.3) on Early Golden Bantam (EGB) maize leaves at 10 days post inoculation (dpi). Disease rates are given as a percentage of the total number of infected plants. Three biological replicates were carried out. n: number of infected maize seedlings. Asterisks above bars indicate significant differences (two-tailed student's t-test). *p*-values are indicated on the bars. **(d)** Disease symptoms caused by *Ustilago maydis* SG200 and SG200Δribo1 mutant strains on Early Golden Bantam (EGB) maize leaves at 12 days post inoculation (dpi). Disease rates are given as a percentage of the total number of infected plants. Three biological replicates were carried out. n: number of infected maize seedlings. Data were presented as mean value ± SD.
